# Loss of muscarinic M5 receptor function manifests disparate impairments in exploratory behavior in male and female mice despite common dopamine regulation

**DOI:** 10.1101/2021.07.11.451965

**Authors:** John A. Razidlo, Skylar M.L. Fausner, Liuchang C. Wang, Salahudeen A. Mirza, Veronica A. Alvarez, Julia C. Lemos

**Author notes:** Corresponding Author: Julia C. Lemos.

## Abstract

There are five cloned muscarinic acetylcholine receptors (M1-M5). Of these, the muscarinic type 5 receptor (M5) is the only one localized to dopamine neurons in the ventral tegmental area and substantia nigra. Unlike M1-M4, the M5 receptor has relatively restricted expression in the brain, making it an attractive therapeutic target. Here we performed an in-depth characterization of M5-dependent potentiation of dopamine transmission in the nucleus accumbens and accompanying exploratory behaviors in male and female mice. We show that M5 receptors potentiate dopamine transmission by acting directly on the terminals within the nucleus accumbens. Using the agonist oxotremorine, we revealed a unique concentration response curve and a sensitivity to repeated stressor exposure. We found that constitutive deletion of M5 receptors reduced exploration of the center of an open field while at the same time impairing normal habituation only in male mice. In addition, M5 deletion reduced exploration of salient stimuli, especially under conditions of high novelty, yet had no effect on hedonia. We conclude that M5 receptors are critical for both engaging with the environment and updating behavioral output in responses to the environment cues, specifically in male mice. A cardinal feature of mood and anxiety disorders is a withdrawal from the environment. These data indicate that boosting M5 receptor activity may be a useful therapeutic target for ameliorating these symptoms of depression and anxiety.

**Significance Statement:** The basic physiological and behavioral functions of the muscarinic M5 receptor remain understudied. Furthermore, its presence on dopamine neurons, relatively restricted expression in the brain, and recent crystallization make it an attractive target for therapeutic development. Yet, most preclinical studies of M5 receptor function have primarily focused on substance use disorders in male rodents. Here we characterized the role of M5 receptors in potentiating dopamine transmission in the nucleus accumbens, finding impaired functioning after stress exposure. Furthermore, we show that M5 receptors can modulate exploratory behavior in a sex-specific manner, without impacting hedonic behavior. These findings further illustrate the therapeutic potential of the M5 receptor, warranting further research in the context of treating mood disorders.

## Introduction

Disruption in both dopamine and acetylcholine transmission has been implicated in preclinical models of depression and anxiety as well as in individuals with mood and anxiety disorders (Belujon and Grace, 2017; Higley and Picciotto, 2014; Nutt et al., 1998; Small et al., 2016). In rodents, genetic, viral or chemogenetic manipulations that reduce the firing of cholinergic interneurons in the nucleus accumbens (NAc) produce depression-like behavior (Cheng et al., 2019; Hanada et al., 2018; Warner-Schmidt et al., 2012). Likewise, manipulations that have inhibited or disrupted normal dopamine transmission in the NAc produce similar depression-like behavior (LeBlanc et al., 2020; Tye et al., 2013).

Acetylcholine and dopamine have a reciprocal modulatory relationship that has been studied widely in the literature. It has been shown that dopamine can induce transient inhibition or pauses in striatal cholinergic interneuron firing via activation of dopamine D2 receptors localized to cholinergic interneurons (Chuhma et al., 2014; Ding et al., 2010; Helseth et al., 2021). Acetylcholine regulation of dopamine signaling is more complex. Focusing on the NAc, it has been demonstrated that synchronous release of acetylcholine via electrical or optogenetic stimulation of cholinergic interneurons can trigger the release of dopamine by activating β2-containing nicotinic acetylcholine receptors (nACh-R) on *en passante* synaptic boutons from dopamine axons within the striatum (Cachope et al., 2012; Rice and Cragg, 2004; Threlfell et al., 2012; Zhou et al., 2001). Acetylcholine also modulates dopamine release and uptake through activation of both Gi/o-coupled and Gq-coupled muscarinic receptors. Activation of M2/M4 autoreceptors can suppress dopamine transmission, likely by suppressing ACh release and the amount of β2-nACh-R activation on dopamine terminals (Threlfell et al., 2010; Threlfell et al., 2012).

Strong evidence points to a role of muscarinic M5 receptors in potentiating dopamine transmission in the ventral striatum (Shin et al., 2015; Yamada et al., 2003). The mechanisms underlying this potentiation are not fully understood, but they likely to involve both an increase in release probability of dopamine as well as suppression of dopamine reuptake, since M5 activation results in a slowing of the decay kinetics of dopamine transients (Shin et al., 2015; Underhill and Amara, 2021). Endogenous release of ACh was shown to potentiate dopamine transmission via these mechanisms as findings showed that ambinomium, an inhibitor of acetylcholinesterase, mimics the potentiation of dopamine transmission observed by muscarinic receptor agonists (Shin et al., 2015). A recent study further demonstrated that the stress-associated neuropeptide corticotropin releasing factor (CRF) increases cholinergic interneuron firing, which in turn, potentiates dopamine transmission via activation of M5 receptors, again indicating that endogenous ACh can also activate M5 receptors to potentiate dopamine transmission (Lemos et al., 2019).

Reduction or ablation of M5 receptor function leads to reduced stimulant-induced psychomotor activation, drug seeking and drug taking (Basile et al., 2002; Fink-Jensen et al., 2003; Gould et al., 2019; Gunter et al., 2018; Thomsen et al., 2005). This appears to be the case for psychostimulants, alcohol, and opioids, revealing a broad impact of these muscarinic receptors as therapeutic target for substance use disorders. Furthermore, suppression of M5 function in the VTA disrupted food reward reinforcement behavior (Wang et al., 2008; Yeomans et al., 2000). These findings point to a potential role of disruption of M5 receptor signaling in depression-like or anxiety-like states, which far fewer studies have investigated. In this study, we examine the impact of M5 receptor deletion on exploratory and hedonic behaviors, two behaviors classically used to assess the expression of anxiety-like or depression-like behaviors in male and female mice.

Here we examined the M5-dependent potentiation of dopamine transmission within the NAc that takes place independent of actions of M5 at the somata of DA neurons, in both male and female mice and across a range of concentrations of the non-selective muscarinic agonist oxotremorine (OXO-M) applied under conditions that favor M5 activation (i.e. in the presence of DhβE). Our findings show that M5-dependent potentiation of dopamine transmission in the NAc is similar between males and females. Interestingly, the concentration-dependence of the M5 response was atypical in that increasing concentrations of OXO-M increased the rate at which a maximal effect was achieved, but not the size of the maximal effect. Moreover, increasing concentrations of OXO-M produced a secondary inhibition that may be due to recruitment of other receptors or desensitization of the M5 receptor. Behaviorally, M5 deletion disrupts normal exploratory behavior in a context-specific fashion without impacting hedonia. This study points to a novel and critical function of M5 receptor activation that we believe will contribute to development of M5 receptor targeted therapeutics.

## Methods

### Animals

All procedures were performed in accordance with guidelines from the Institutional Animal Care and Use Committee at the University of Minnesota. Male and female mice (p60-180) were group housed and kept under a 12h-light cycle (6:00 ON/18:00 OFF) with food and water available *ad libitum*. Breeding was done under “summer photoperiod” (14:10 light cycle) and then post-weaning, animals were transferred to our Investigator Management Housing Area and kept under a 12:12 light cycle for at least a week before use. M5 -/- (knock-out) mouse line was a generous gift of Dr. Jürgen Wess. M5 wild-type (WT, M5+/+), heterozygous (HET, M5+/-) and knock-out (KO, M5-/-) littermates were generated using a M5 HET x M5 HET breeding scheme that produced approximately 25% WT, 50% HET, 25% KO. The M5 HET x M5 HET (C57BL/6J background) breeding strategy was employed to ensure uniform maternal care during development. However, a caveat to this strategy is that there is a non-uniform number of WT, HET, KO mice in each cage. We found that this disrupted normal social interaction behavior and therefore excluded these experiments from our study. The low yield of WT and KO mice also made it prohibitive to parse estrous-cycle dependent effects, though this is of interest for future studies.

For neuroanatomy and behavior experiments, we used M5 WT littermates only. For voltammetry experiments, it was not possible to exclusively use M5 WT littermates given the small percentage produced. Thus, we combined M5 WT littermates with C57BL/6J mice bred inhouse under identical conditions. We found no difference between these groups and therefore pooled the data.

Genotyping was outsourced to Transnetyx ®. Original development of M5 KO mice was completed through insertion of a neomycin-resistance cassette (*NRC*) replacing the start of *Chrm5* gene (first 750 base pairs (b.p.) (first 250 amino acids including the translation start site). (Yamada et al., 2001). Mice were classified as WT if they were positive for *Chrm5* and negative for *NRC*, HET mice were positive for both *Chrm5* and *NRC*, and KO mice were negative for *Chrm5* and positive for *NRC*. (*Chrm5*: Forward: CCATCACAAGACCACTGACATACC; Reverse: GCCATGCCAAGCCGATCA; *NRC*: Forward: GGGCGCCCGGTTCTT; Reverse: CCTCGTCCTGCAGTTCATTCA). Genotyping was performed on tail-tip samples collected on the day of weaning (p21).

### Fluorescent In Situ Hybridization (ISH) using RNAscope ®

Brains were rapidly dissected, and flash frozen in isopentane on dry ice. Brains were kept in a −80°C freezer until they were sectioned. Coronal sections (16 μm) containing the VTA were thaw mounted onto Superfrost plus slides (Electron Microscopy Sciences) utilizing a Leica CM 1860 cryostat maintained at −20°C. Prior to sectioning, brains were equilibrated in the cryostat for at least 2 hrs (overnight is optimal). Slides were cleaned with RNAseZap® RNAse Decontamination Solution to prevent mRNA degradation. Slides were stored at −80°C. RNAscope® ISH was conducted according to the Advanced Cell Diagnostics (ACD) user manual. Briefly, slides were fixed in 10% neutral buffered formalin for 20 min at 4°C. Slides were washed 2×1 min with 1x PBS, before dehydration with 50% ethanol (1 x 5 min), 70% ethanol (1 x 5 min), and 100% ethanol (2 x 5 min). Slides were incubated in 100% ethanol at −20°C overnight. The following day, slides were dried at room temperature (RT) for 10 min. A hydrophobic barrier was drawn around the sections using a hydrophobic pen and allowed to dry for 15 min at RT. Sections were then incubated with Protease Pretreat-4 solution for 20 min at RT. Slides were washed with ddH2O (2 x 1 min), before being incubated with the appropriate probes for 2 hr at 40°C in the HybEZ oven (ACD). Our preliminary data demonstrated that the commercially available ACD probe for *Chrm5* (ACD Cat. #: 495301) had an unacceptable amount of signal in the M5 KO mouse. Thus, we collaborated with ACD to design a custom *Chrm5* probe directed at the first 750 bp of the *Chrm5* gene which has been replaced with an NRC in the M5 KO (ACD Cat. #: 1052411, *Mm-Chrm5-O2*). This probe was multiplexed with the probe directed at *tyrosine hydroxylase (Th)* (ACD Cat. # 317621). Following incubation with the appropriate probes, slides were subjected to a series of amplification steps at 40°C in the HybEZ oven with 2 x 2 min washes (with agitation) in between each amplification step at RT. Amplification steps were as followed: Amp 1 at 40°C for 30 min. Amp 2 at 40°C for 15 min. Amp 3 at 40°C for 30 min. Amp 4-Alt B at 40°C for 15 min. A DAPI-containing solution was applied to sections (one slide at a time) at RT for 20 sec. Finally, slides were cover-slipped using ProLong Gold Antifade mounting media (Invitrogen) and stored at 4°C until imaging.

### Image analysis and quantification for RNAscope®

Sections were imaged using Keyence BZ-X710 epifluorescent microscope and corresponding software. Unique 20x images of the VTA were acquired from M5 WT male and female littermates using the same software and hardware settings. The settings were titrated for each specific experimental probe. Quantification was done using Fiji/ImageJ software. Numbers of DAPI and TH+ cells were automatically generated using the particle counter function in ImageJ. A TH+ cytosolic mask was generated by subtracting a DAPI+ mask from the original TH+ mask. This TH+ cytosolic mask was then used to assess M5/TH co-expression within the VTA and puncta number. Thresholding was kept consistent across images.

### Fast Scan Cyclic Voltammetry (FSCV)

Coronal slices (240 μm) containing NAc core were prepared from M5 WT, M5 HET or M5 KO mice. Slices were cut in ice-cold cutting solution (in mM): 225 sucrose, 13.9 NaCl, 26.2 NaHCO_3_, 1 NaH_2_PO_4_, 1.25 glucose, 2.5 KCl, 0.1 CaCl_2_, 4.9 MgCl_2_, and 3 kynurenic acid. Slices were maintained in oxygenated artificial cerebral spinal fluid (ACSF) containing (in mM): 124 NaCl, 2.5 KCl, 2.5 CaCl_2_, 1.3 MgCl_2_, 26.2 NaHCO_3_, 1 NaH_2_PO_4_, and 20 glucose (~310-315 mOsm) at RT following a 1hr recovery period at 33°C. Carbon fiber (7 μm diam., Goodfellow) electrodes were fabricated with glass capillary (602000, A-M Systems) using a Sutter P-97 puller and fiber tips were hand cut to 100-150 μm past the capillary tip. Immediately prior to the experiments they were filled with 1M KCl internal solution. Recordings were conducted at 31-33°C maintained with a Warner inline heating system. The carbon-fiber electrode was held at −0.4 V and a voltage ramp to and from 1.3 V vs. Ag/AgCl (400V/s) was delivered every 100 ms (10 Hz). Before recording, electrodes were conditioned by running the ramp at 60 Hz for 10 min and at 10 Hz for another 10 min. Dopamine transients were evoked by electrical stimulation delivered through a glass microelectrode filled with ACSF. A single monophasic pulse (4 msec, 300 μA) was delivered to the slice in the absence or presence of the nACh-R antagonist dihydro-β-erythroidine (DhβE, 1 μM). Data was acquired with using a Dagan headstage and amplifier and National Instruments PCI boards. Data was acquired and analyzed using Demon Voltammetry and Analysis software package (Yorgason et al., 2011). Experiments were rejected when the evoked current did not have the characteristic electrochemical signature of dopamine assessed by a current-voltage plot.

### Behavior

Prior to the start of behavioral experiments, mice were handled and acclimated to the testing room for five days. Behavioral testing chambers for open field, novel object or food exploration, light/dark box and elevated zero maze were contained in individual custom built sound attenuated chambers that were equipped with overhead light (with dimmer switch) and white noise fan. Mice were video monitored during behavioral testing and data was acquired and analyzed using Noldus Ethovision (v. 14,15) software. *Open field (OF), novel object exploration, novel palatable food exploration*: Animals were placed in a circular arena (50 cm diameter, 40 cm height) for 60 minutes on day 1. On day 2, mice were placed in the OF chamber and allowed to habituate for 60 minutes. Mice were then placed back in their home cage (HC) for 5 minutes. A novel object (NO;5 cm cylindrical bottle cap from a 1L pyrex laboratory bottle) or a container holding novel palatable food (NF; bacon softies covered with a fine mesh, BioServ, F3580) was placed in the center of the arena in a counterbalanced fashion. Mice were allowed to explore for 30 minutes. Mice were then taken out of the arena and placed back in HC for 5 minutes. Subsequently, the alternate stimulus was placed in the center of the arena and mice were allowed to explore for another 30 minutes. *Elevated zero maze*: The elevated zero maze (EZM) customized for mice was purchased from Med Associates (64 cm diameter, 65 cm height). The EZM purchased came with hinged covers for the closed compartments, giving the ability to have the closed compartments fully enclosed or have the top of the closed compartment exposed. We used two separate conditions for our studies. In one condition, the EZM was illuminated by bright light (161 lux) and the closed compartments were fully enclosed. In the second condition, mice experienced the EZM under dim conditions (49 lux) with the tops of closed compartments exposed. Mice were exposed to both conditions in a counterbalanced fashion and were exposed to each condition 7 days apart. The prediction was that mice should modulate their behavior, specifically their time and entries into the open compartments based on which condition they were exposed to. Each EZM session was 5 minutes long. *Sucrose Preference Test*: Mice were individually housed for these experiments. Mice were acclimated to the two-bottle home cage apparatus, in which both bottles contained water, for three days. On the fourth day, one bottle contained 4% sucrose and one contained water. The side in which the sucrose water was placed was counterbalanced. Sucrose and water bottles were weighed for three days, alternating sides each day, and results were averaged. *Light-dark box*: A separate cohort of mice were used for these experiments that had no prior experience with other behavioral chambers. Mice were placed in the dark compartment of the light-dark box with bright overhead illumination (161 lux) (Stoelting, Cat. 10-000-355). Activity and time allocation in the light or dark compartment was assessed over a 5 min duration. Mice were subsequently placed back in HC for 5 min. Mice were then reintroduced to the testing chamber (starting in the dark side) with a novel palatable food placed in the bright side. Activity and time allocation in the light or dark compartment was assessed over a 5 min duration.

### Stressor Exposure: Repeated Forced Swim Stress

On day 1, male mice were placed in a 4L bucket containing 30 ± 1°C water for 15 minutes. On day 2-7, mice were re-exposed to the 30 ± 1°C water for 5 minutes at different times of day across each day. Mice demonstrated robust increase in immobility over time (data not shown). Mice were prepared for fcsv experiments 24-72 hrs after the last swim exposure. *Chronic Restraint Stress*: Mice were confined to a mouse restrainer (50 mL conical tube with air holes) for a period of 30 minutes, every day at 10 a.m. for two weeks. For both paradigms, control mice were handled for the same number of days as the corresponding stressor exposure.

### Statistics

Statistical analysis was performed in Prism (GraphPad) and Excel. Two-tailed unpaired t-test, two-tailed paired t-test, 1W ANOVAs or 2W Repeated Measures (2WRM) ANOVAs were used when appropriate and are stated in the results. 1W and 2W ANOVAs were followed up with a Dunnett’s or Sidak-corrected t-test comparisons. All data are presented as mean ± SEM. Results were considered significant at an alpha of 0.05. * = t-tests or post-hoc t-tests following 1W or 2W ANOVAs, # = significant interaction or second post-hoc t-test. & = 2W ANOVA, main effect, δ = trend 0.100 >p>0.05. * p < 0.05, ** p < 0.01, *** p < 0.001, **** p < 0.0001.

## Results

### Chrm5 is ubiquitously expressed on Th+ neurons in the VTA

The expression of *Chrm5* mRNA that encodes the M5 receptor protein was assessed. According to the Allen Brain Atlas *in situ* hybridization data, *Chrm5* has a markedly limited expression compared to the other four cloned muscarinic acetylcholine receptors *Chrm1-4* (www.alleninstitute.org) (Figure 1a) (Lein et al., 2007). One concentration of *Chrm5* mRNA is the ventral midbrain (Figure 1a). To assess *Chrm5* mRNA expression specifically on dopamine neurons we used a custom RNAscope® probe targeted to the first 750 b.p. of the *Chrm5* gene where the neomycin-resistant cassette was inserted to generate the M5 KO mouse (see Methods for details). Using this probe, we confirmed that *Chrm5* mRNAs are localized to dopamine neurons in the VTA and substantia nigra *pars compacta* by multiplexing with a probe targeted to *Th* mRNAs. Using an automated methodology in ImageJ, we determined that *Chrm5* is localized to 84% of *Th+* neurons in VTA (Figure 1b,c). When comparing males and females, we found a significant difference in *Chrm5* mRNA puncta in putative dopamine neurons that were isolated using a *Th* mask (with nucleus excluded) in Image J (males: 593 ± 40; females: 756 ± 42 particles, t-test, t_7_ = 2.499, p = 0.041, n =6,3 respectively). We speculated that the higher expression of *Chrm5* mRNA in females may manifest on functional differences, which motivated the subsequent analysis. We began by confirmation and validation of previous reports that M5 receptor activation potentiated dopamine transmission in the NAc.

**Figure 1.**
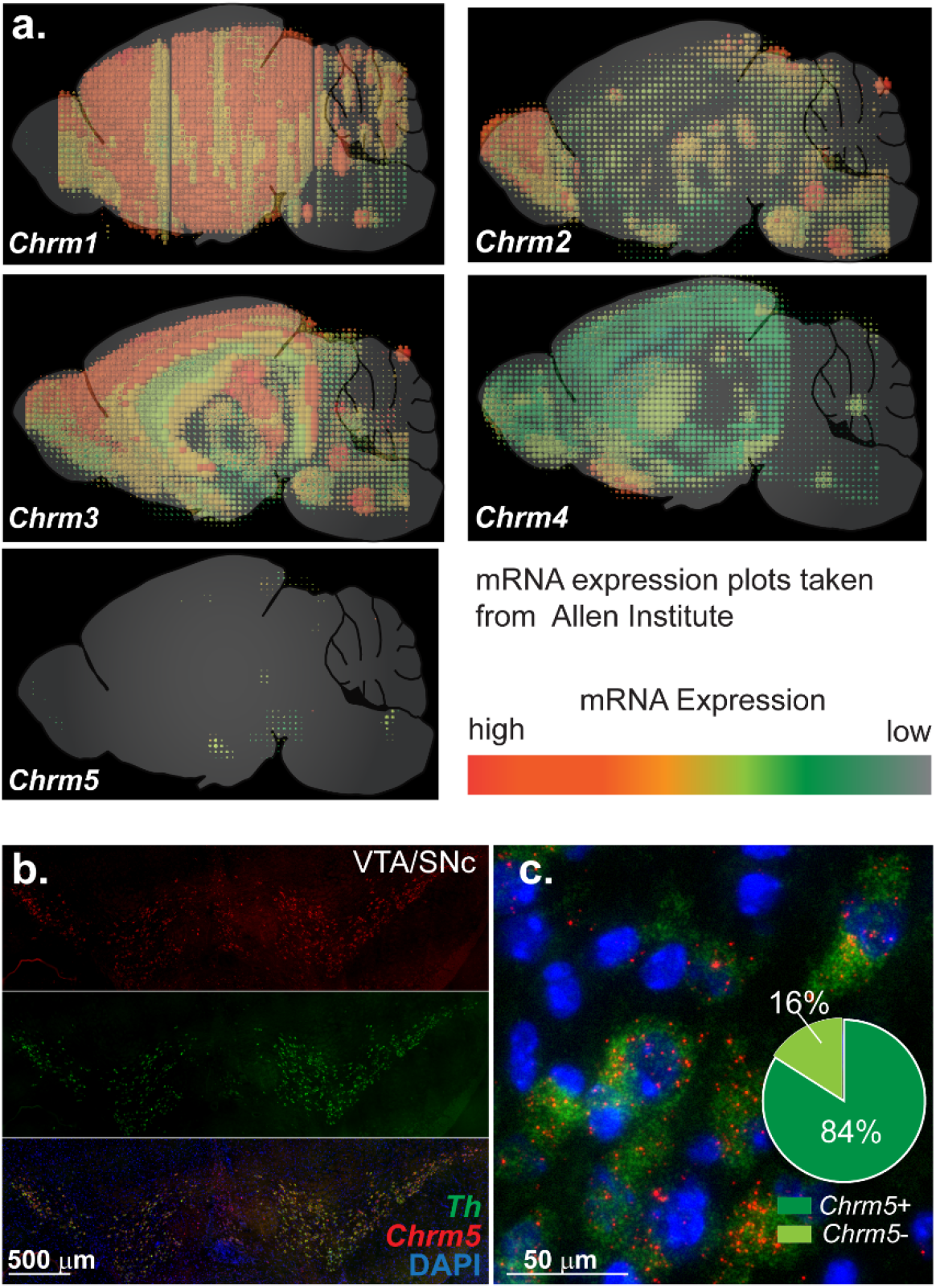
Muscarinic M5 receptors are localized to midbrain dopamine neurons. a) Composite images of brain-wide mRNA expression data for *Chrm1-5* (M1-M5) based on *in situ* hybridization data collected by the Allen Institute and made available on www.alleninstitute.org. b,c) Low power (b) and high power (c) fluorescent images of *Th* (green) and *Chrm5* (red) mRNA expressed in the VTA. Quantification of *Chrm5/Th* co-expression in the VTA of male and female M5 WT mice, 22,177 *Th+* cells counted from 9 animals (6 males, 3 females).

### M5-dependent potentiation of dopamine transmission

The reciprocal interaction between cholinergic and dopaminergic systems within the striatum is complex. Electrical stimulation within the NAc core produces a composite dopamine transient, a summation of direct dopamine fiber stimulation and indirect dopamine release. Indirect dopamine sources arise from stimulation of cholinergic interneurons and acetylcholine release that mediate dopamine transmission via activation of β2-containing nACh-Rs on dopamine varicosities. (Figure 2-1b). We first sought to identify any sex or genotype differences in the nACh-R mediated component of the dopamine transient, which is sensitive to the blocker DhβE. DhβE (1μM) produced a 56% reduction in the peak dopamine transient in males (Figure 2-1, n =13) and 52% reduction in females (n = 31, not shown). Interestingly, in knockout mice for the muscarinic M5 receptor (M5 KO), dopamine transients were less sensitive to the nicotinic blocker (46 ± 1 % reduction from baseline in male M5 KO compared to 56 ± 2 in male wildtype (WT) mice; t-test, t_18_ = 3.198, p = 0.0050, n = 7-13). This is a small quantitative difference; qualitatively, the contribution of nACh-R activation to the electrically evoked composite dopamine transient is similar.

**Figure 2.**
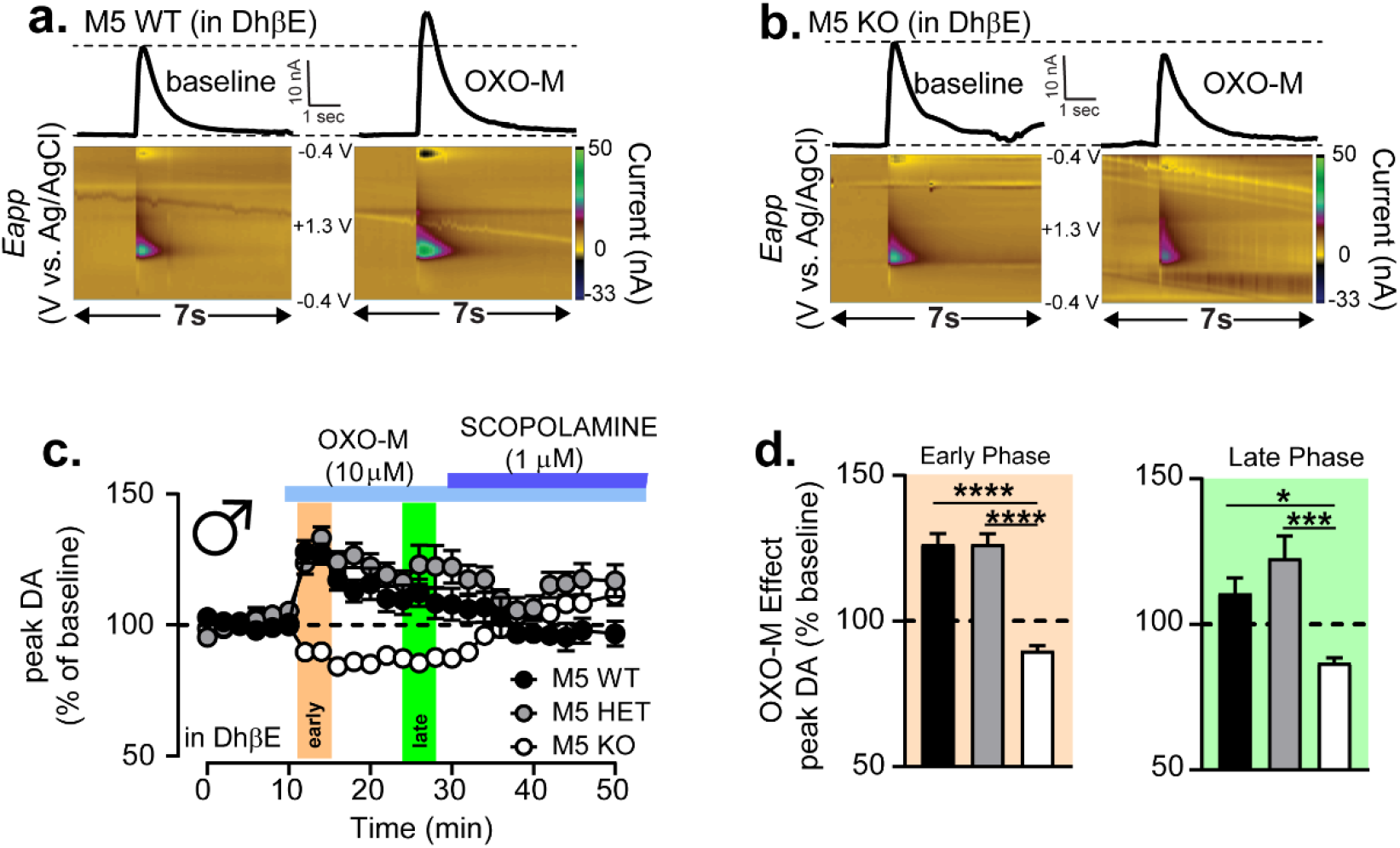
Muscarinic M5 receptor activation potentiates dopamine transmission in the NAc core. a) Representative current x time and 3D fast scan cyclic voltammetry color plots for electrically evoked dopamine transients recorded in M5 WT (a) and M5 KO (b) mice prior to and following OXO-M (10μM) in slices maintained in DhβE (1 μM). c) Time course of normalized peak amplitude of electrically evoked dopamine transients recorded from *ex vivo* NAc slices taken from M5 WT, HET and KO male mice in which OXO-M, followed by Scopolamine, is bath applied to the slice. d) Average OXO-M effect for M5 WT, HET and KO mice in the first (early, left) and last (late, right) four minutes of the OXO-M bath application.

When we assessed the function of the muscarinic autoreceptors M2/M4, which are expressed in the cholinergic interneurons and inhibit ACh transmission, we used the non-selective agonist oxotremorine (OXO-M, 1μM) and tested it in the <u>absence</u> of DhβE. We found that OXO-M effect was nearly identical between M5 WT and M5 KO mice (M5 WT: 59 ± 4; M5 KO: 60 ± 6 % reduction from baseline, t-test, t_7_ = 0.1590, p = 0.8782, n = 4-5). From this, we conclude that the cholinergic mechanisms responsible for triggering dopamine release which involve β2-nACh-Rs and the mechanism responsible for negative feedback on ACh release which involve M2/M4 receptors are for the most part intact in M5 KO mice.

In the presence of DhβE, electrically evoked dopamine transients are derived solely from excitation of dopamine fibers and as such they are not affected by activity of M2/M4 autoreceptors. Under these conditions, the non-selective agonist OXO-M instead potentiates the peak amplitude of the dopamine transient in an M5-dependent fashion (Shin et al., 2015). In this current study, we replicate and extend the findings of Shin et al., 2015 by including M5 HET in our analysis and assessing the temporal kinetics of this response across several OXO-M concentrations in both male and female mice.

In agreement with previous findings, OXO-M (10 μM) produces a significant potentiation of the peak amplitude of the electrically evoked dopamine transient in the presence of DhβE in male WT mice. The potentiation is fully reversed by the non-selective muscarinic antagonist scopolamine (1 μM), confirming that the effect is muscarinic receptor dependent. Interestingly, we found that this concentration of OXO-M produces a rapid increase in dopamine transmission to 126 ± 4% of baseline within the first five minutes. However, this is transient, and it significantly attenuates over time to steady state amplitude of 111± 5% of baseline. Thus, there was a difference in the early vs. late phase of the OXO-M effect in male WT mice (paired t-test, t_40_ = 2.433, p = 0.0195, n = 21, Figure 2a-d). Interestingly, mice with heterozygote deletion of M5 receptor gene (M5 HET) show an intact OXO-M induced potentiation of dopamine transmission that was not different than WT mice (both 126 ± 4% of baseline in early phase). However, unlike WT mice, M5 HET mice displayed no significant attenuation of the OXO-M response over time (M5 HET: early phase: 126 ± 4% of baseline; late phase: 123 ± 8% of baseline, paired t-test, t_12_ = 0.5061, p = 0.6219, n = 13, Figure 2c). We confirmed that this OXO-M induced potentiation of dopamine transmission requires M5 receptor activation using the M5 KO littermates. Indeed, the OXO-M mediated potentiation was absent in M5 KO mice and there was even a small but significant inhibition of the peak dopamine transient during OXO-M application. The inhibition was evident in the early phase and remained stable through the late phase confirming that OXO-M potentiation of dopamine transmission was dependent on M5 receptors under these conditions (1W ANOVA, M5 WT, HET, KO early phase: F_2,44_ = 34.52, p < 0.0001, post-hoc Tukey’s t-tests, p < 0.0001; late phase: F_2,45_ = 8.623, p = 0.007, post-hoc Tukey’s t-tests, M5 KO vs. M5 WT, p = 0.0135, vs. M5 HET, p = 0.0005, n = 13-21, Figure 2c.d).

We were intrigued by the time course of the OXO-M effect on M5-dependent potentiation in control mice and probed this further by examining a range of OXO-M concentrations in both males and females. In all cases, we used scopolamine (1 μM) to reverse the effects of OXO-M and confirm the involvement of muscarinic receptors (Figure 3-1). Increasing concentrations of OXO-M from 0.1 μM to 25 μM (0.1, 0.5, 2.5,10. 25) did <u>not</u> increase the peak DA transients but rather significantly sped up the time to peak across concentrations (Figure 3a-f, Figure 3-1) (Peak OXO-M effect: 2W ANOVA, F_4,71_ = 0.2917, p = 0.8824; Time to peak: 2W ANOVA, main effect of concentration, F_4,71_ = 20.84, p < 0.0001). There was also a significant difference in the late phase attenuation of the response such that at 25μM, the response rapidly returns to baseline levels. This is demonstrated by comparing the early and late phase concentration response curves for males and females (Figure 3d) (male: 2W ANOVA, concentration x time interaction, F_4,40_ = 7.347, p = 0.0002, n = 5-20; female: 2W ANOVA, concentration x time interaction, F_4,32_ = 12.66, p < 0.0001, n = 7-8) and calculating the % attenuation as Early - Late phase (% baseline, Figure 3-1). Males and females had very similar responses to OXO-M across concentrations. The one exception was at 0.5 μM concentration where we did see a significant difference in the overall time course (Figure 3-1) as well as trend in the time to peak (post-hoc t-test, p = 0.0579). However, we conclude that generally there were only slight differences in OXO-M response between males and females.

**Figure 3.**
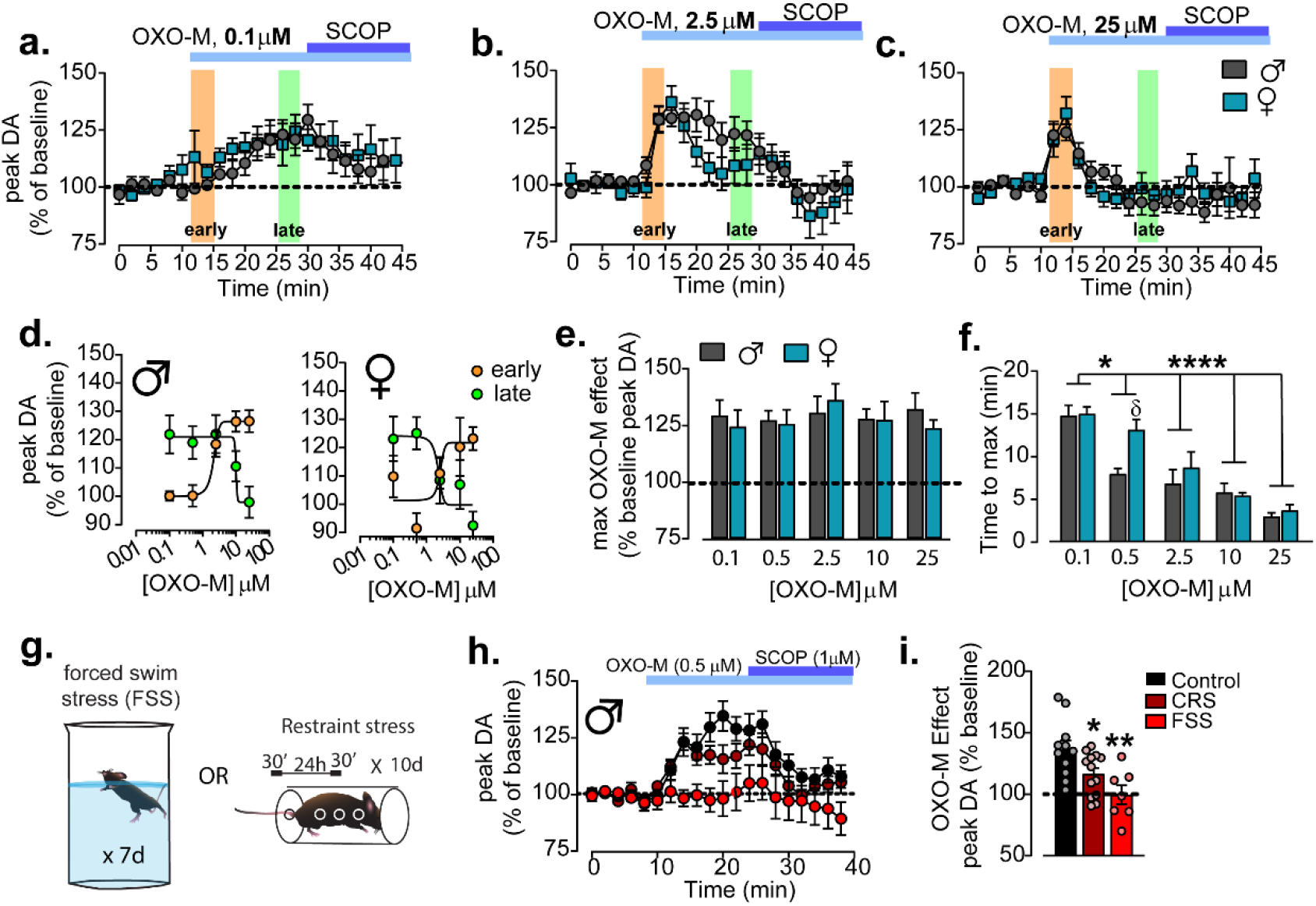
M5-dependent potentiation of dopamine transmission displays unique temporal kinetics and sensitivity to stress. a-c) Time course of normalized peak amplitude of electrically evoked dopamine transients recorded from *ex vivo* NAc slices taken from male (grey) and female (teal) M5 WT mice in which OXO-M (0.1 μM, 2.5 μM, 25 μM) followed by scopolamine (1μM) is bath applied to the slice. d) OXO-M concentration response curve for the early (orange) and late (green) stages of OXO-M bath application for males (left) and females (right) e) Peak OXO-M effect 0.1, 0.5, 2.5, 5, 10 and 25 μM OXO-M in males and females. f) Time to peak for 0.1, 0.5, 2.5, 5, 10 and 25 μM OXO-M in males and females. g) Schematic demonstrating stress paradigms used for h. and i. h) Time course of normalized peak amplitude of electrically evoked dopamine transients recorded from *ex vivo* NAc slices taken from stress-naïve control male mice and male mice exposed to repeated forced swim stress for 7 days or chronic restraint stress for 10 days in which OXO-M (0.5 μM), followed by Scopolamine is bath applied to the slice. i) Mean OXO-M (0.5 μM) response in control, forced swim stress (FSS) and chronic restraint stress (CRS) exposed mice.

We previously showed that repeated exposure to stress disrupts CRF-mediated potentiation of dopamine transmission in the NAc core (Lemos et al., 2012). Here, we further assessed whether repeated stress similarly disrupted M5 modulation of dopamine transmission. We exposed mice to either 7 days of repeated swim stress or 10 days of repeated restraint stress. In both cases, stress reduced the maximal potentiation induced by 0.5 μM OXO-M compared to stress-naive mice (Figure 3g-i) (stress-naïve: 137 ± 7, CRS: 117 ± 5, FSS: 100 ± 8 % baseline, 1W ANOVA, F_2,28_ = 7.325, p = 0.0028, Dunnett’s t-test: control vs. CRS, p = 0.0427, vs. FSS, p = 0.0016, n = 7-13).

### M5 deletion manifest disparate behavioral deficits in the open field in male and female mice

Based on our finding that stress disrupts M5 signaling, we wondered how genetic constitutive M5 deletion would impact dopamine-dependent behaviors including locomotion, exploratory behaviors, and hedonic behaviors (Duzel et al., 2010). We first assessed the behavior of M5 WT, HET and KO male and female littermates in the novel open field task performed under dim lighting conditions (49 lux). Generally, females showed enhanced locomotion in the open field compared to males (2W-ANOVA, main effect of sex, F_1,108_ = 12.14, p = 0.007). We found no difference in spontaneous horizontal locomotion across genotypes for neither males nor females (Figure 4a,b,c,d,f,g) (males: 1W ANOVA on total time, F_2,66_ = 1.252, p = 0.2925, n = 19-31; females, 1W ANOVA on total time, F_2,42_ = 0.6094, p = 0.5484, n = 14-16). In addition to horizontal ambulatory behavior, we also analyzed rearing behavior during the first 10 minutes of the open field for M5 WT and KO male and female littermates as it has been shown to be a secondary measure of exploration, indicative of vigilance and dependent on dopamine transmission (Sturman et al., 2018; Vallone et al., 2002). While there was no difference in number of unsupported rears between male mice, there was a significant reduction in rearing behavior in female M5 KO compared to WT littermates (male: WT: 7 ± 1, KO: 8± 2 rears, t-test, t = 0.0356, p = 0.7103, n = 14 each; female: WT: 14 ± 2, KO: 9 ± 1 rears, t-test, t = 2.338, p = 0.0265, n = 15-16). We assessed time in center and found that M5 deletion significantly reduced time in center during the first 30 minutes only in male mice. This does not occur in female M5 KO mice which were similar to WT and HET. In both male and female M5 WT mice, exploration of the center of the open field is always larger in the first 30 mins than in the last 30 mins, as expected for exploratory behavior that extinguishes as time passes (Fig. 4i). However, in both male and female M5 KOs this habituation in exploration of the center is disrupted. M5 HET mice had an intermediate phenotype, where males showed similar behavior to control M5 WT, while M5 HET females showed disruption in the habituation, similar behavior to M5 KO (Figure 4i,j)(Males, M5 WT,HET,KO: 2W ANOVA, time x genotype interaction, F_2,61_ = 5.342, p = 0.0073, post-hoc Sidak’s t-test, first 30 minutes for WT vs. HET, p = 0.8945, WT vs KO, p = 0.0299, HET vs. KO, p = 0.0477, post-hoc t-test of 0-30 vs. 30-60 minutes, WT = p = 0.0208, HET = p < 0.0001, KO = p = 0.9964, n = 14-25; Females: M5 WT,HET,KO: 2W ANOVA, main effect of time, F_1,40_ = 18.88, p 0.0001, post-hoc Sidak’s t-test, first 30 minutes for WT vs. HET, p = 0.6723, WT vs KO, p = 0.6466, HET vs. KO, p > 0.9999, post-hoc t-test of 0-30 vs. 30-60 minutes, WT = p = 0.0064, HET = p = 0.1270, KO = p = 0.0894, n = 13-16). Under these conditions, females spent more time in the center compartment compared to males, however, there was not a significant sex x genotype interaction (2W ANOVA, main effect of sex, F_1,101_ = 6.925, p = 0.0098, sex x genotype interaction, F_2,101_ = 0.6531, p = 0.5226). Together, these behavioral findings point to deficit in exploratory behavior in mice lacking muscarinic M5 receptors. We also found sex specific differentiation on the impact of M5 deletion on exploration, indicating that M5 function regulates different aspects of these behaviors.

**Figure 4.**
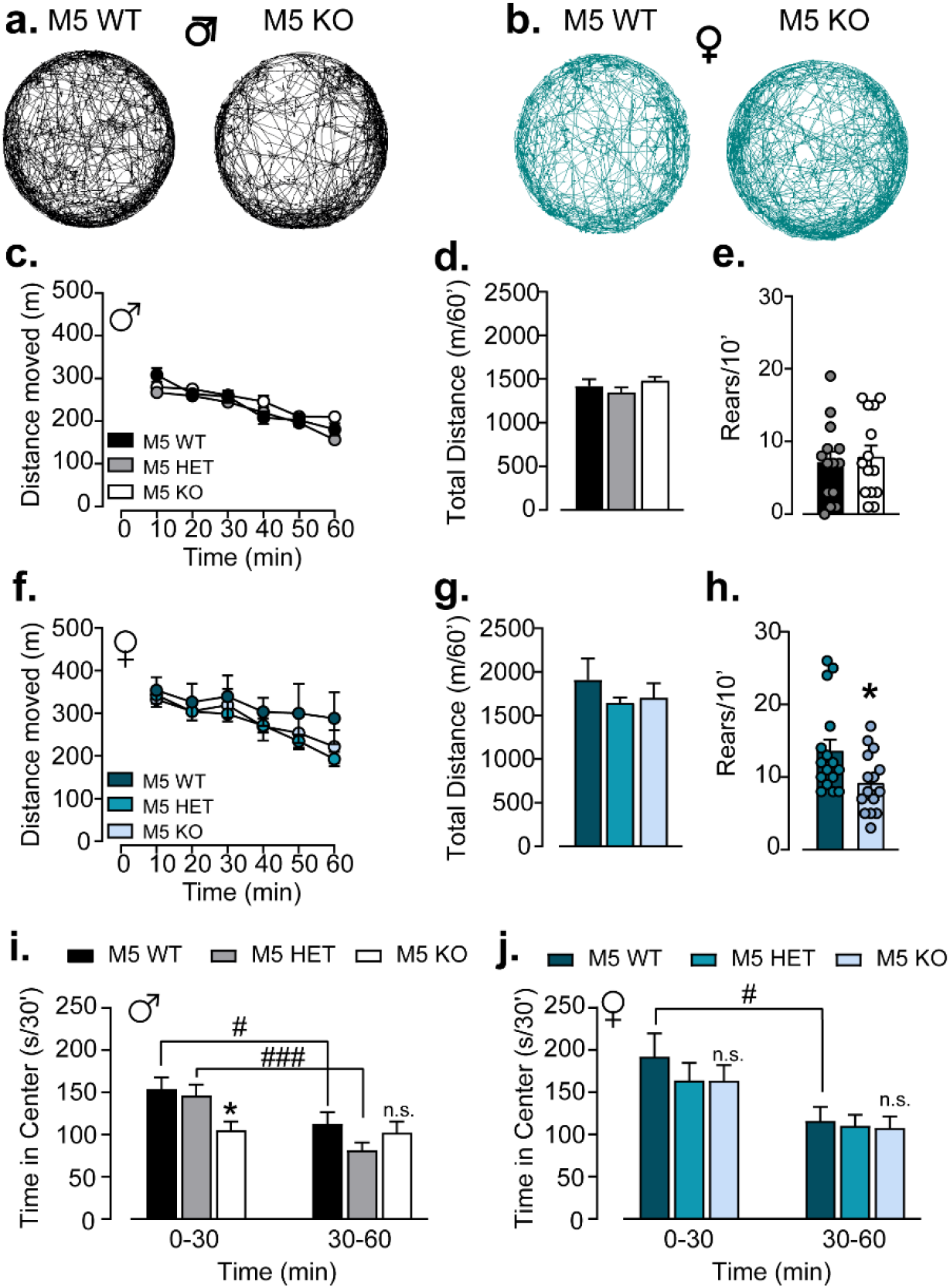
M5 deletion produces disparate behavioral deficits in the open field in males and females. a,b) Representative motor tracks extracted from video monitoring of the mice in the open field for male (a) and female (b) M5 WT (left) and M5 KO (right) mice. c) Time course of spontaneous locomotor activity during 60 min exposure to novel open field for male M5 WT, HET, KO. d) Mean total motor activity in 60 min novel open field session for male M5 WT, HET, KO e) Total number of unsupported rears in the first 10 mins of novel open field session for male M5 WT and KO. f) Time course of spontaneous locomotor activity during 60 min exposure to novel open field for female M5 WT, HET, KO. g) Mean total motor activity in 60 min novel open field session for female M5 WT, HET, KO h) Total number of unsupported rears in the first 10 mins of novel open field session for female M5 WT and KO. i) Time in center of novel open field in the first and second 30 min of a 60 min open field session for male M5 WT, HET and KO mice. j) Time in center of novel open field in the first and second 30 min of a 60 min open field session for female M5 WT, HET and KO mice.

On the subsequent day, we used the same open field apparatus to assay the response to novel stimuli. Mice were habituated to the OF for 60 min under dim conditions. We investigated time spent exploring both a novel object (1L bottle cap) or novel palatable food (bacon softies). The novel food was contained in a dish, covered by a mesh such that mice could smell the food but not consume it. The novel object (NO) and novel palatable food (NF) exposure were counterbalanced across two sessions. We had hypothesized that mice would increase their exploration time toward NF compared to NO in WT, but not KO mice. While there was a trend for this pattern in males, overall, there were no significant differences in exploration of the two stimuli in either WT or KO male or female mice. There was, however, an overall main effect of genotype on exploration in males only (Figure 5a,b,d) (Males: 2W ANOVA, stimuli x genotype interaction, F_1,32_ = 0.6007, p = 0.4440, main effect of genotype, F_1,32_ = 4.627, p = 0.0391, n = 17 each, Females: 2W ANOVA, stimuli x genotype interaction, F_1,26_ = 0.6583, p = 0.4245, main effect of genotype, F_1,26_ = 0.9796, p = 0.3314, n = 13-15). We then analyzed the exploratory response to the first stimulus compared to the second stimulus exposure regardless of stimulus type. In this case, there was a significant main effect of both genotype and time, indicating that male M5 KO mice have a deficit in exploratory behavior, particularly during the first stimulus presentation. In contrast to our findings in males, females had no significant differences in exploratory behavior (Figure 5c,e) (Males: 2W ANOVA, main effect of time, F_1,32_ = 28.44, p < 0.0001, main effect of genotype, F_1,32_ = 4.627, p = 0.0391, post-hoc t-test of M5 WT vs. KO for stimulus 1, p = 0.0259, n = 17 each, Females: 2W ANOVA, main effect of time, F_1,22_ = 1.713, p = 0.2041, main effect of genotype, F_1,26_ = 0.4313, p = 0.5181, n = 11-13).

**Figure 5.**
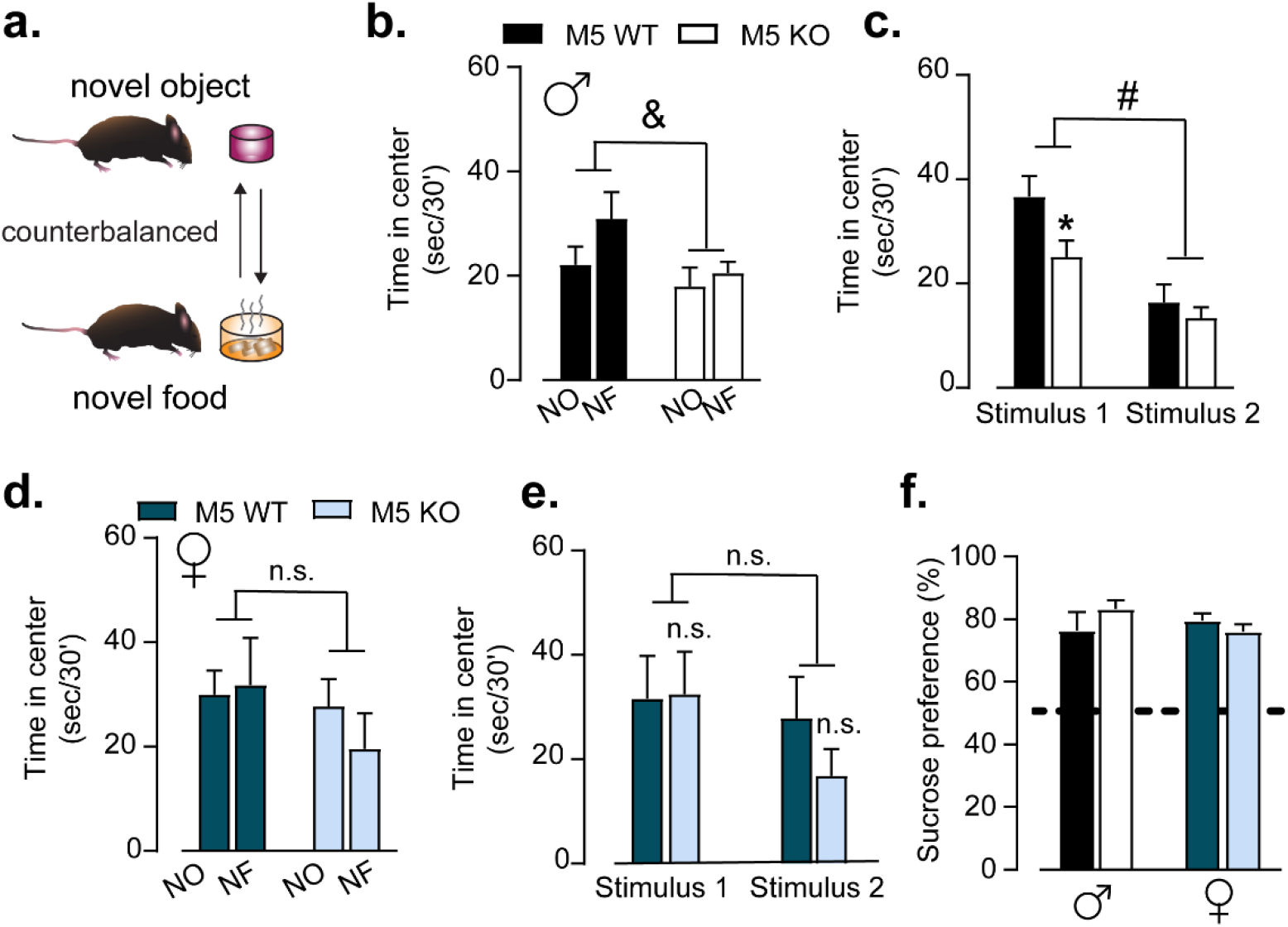
M5 deletion reduces novelty exploration only in males without impacting hedonia. a) Schematic diagram of novel stimulus paradigm. b) Time in center during a 30 min session in which either a novel object (NO) or novel palatable food (NF) was placed in the center of an open field for male M5 WT or M5 KO. c) Time in center during a 30 min session for Stimulus 1 (presented first) or Stimulus 2 (presented second) regardless of stimulus type placed in the center of an open field for male M5 WT or M5 KO. d) Time in center during a 30 min session in which either a novel object (NO) or novel palatable food (NF) was placed in the center of an open field for female M5 WT or M5 KO. e) Time in center during a 30 min session for Stimulus 1 (presented first) or Stimulus 2 (presented second) regardless of stimulus type placed in the center of an open field for female M5 WT or M5 KO. f) Sucrose preference (4% sucrose) for male or female M5 WT or KO. Sucrose preference = sucrose volume consumed/sucrose + water volume consumed x 100.

It has been demonstrated that the non-selective antagonist scopolamine reduces sucrose preference, a test of anhedonia, when administered systemically. Since M5 KO mice, males in particular, display a reduction of exploration of appetitive stimuli, we wondered if these same mice displayed anhedonia. However, we found that neither male nor female M5 KO mice displayed a deficit in sucrose preference (using 4% sucrose solution) compared to M5 WT littermate controls (Figure 5f) (2W ANOVA, all comparisons, p > 0.05).

### M5 deletion disrupts normal behavioral adaptations to changing environmental conditions

Time allocation in the center of an open field is sensitive to anti-anxiety drugs, and thus is considered a test of anxiety-like behavior (Prut and Belzung, 2003). We used the elevated zero maze as a secondary measure of anxiety-like behavior. Performance in the elevated zero maze (EZM) and the more commonly used elevated plus maze (EPM) are sensitive to changes in the environmental conditions, especially light (Walf and Frye, 2007). Mice were exposed to the EZM under two different, counterbalanced conditions: dim light, where the “closed” arms are not covered, and bright light, where the “closed” arms are covered (Figure 6a). It has been shown that modulating the luminescence as well as conditions of the closed arms (i.e. transparent vs. opaque walls) impacts time in open arms. Generally, there was no overall effect on time in open arms by genotype. However, M5 WT male mice spent significantly more time in the open arms under the dim conditions, as did M5 HET. However, M5 KO mice did not significantly change their behavioral response to EZM between conditions (Figure 5b, 2W ANOVA, main effect of lighting, F_1,100_ = 16.48, p < 0.0001, main effect of genotype, F_2,100_ = 3.521, p = 0.0333, post-hoc t-test, bright vs. dim, M5 WT: p = 0.0391, M5 HET: p = 0.0008, M5 KO: p = 0.5964, n = 13-25). While females showed the same qualitative pattern, the effects were not statistically significant for any comparison. M5 WT females showed a trend in difference between dim and bright conditions that was absent in M5 HET and KO mice. However, overall, the quantitative differences were not significant (Figure 5c, 2W ANOVA, main effect of lighting, F_1,47_ =6.705, p = 0.0128, main effect of genotype, F_2,47_ = 0.8458, p = 0.4356, post-hoc t-test, bright vs. dim, M5 WT: p = 0.0852, M5 HET: p = 0.8487, M5 KO: p = 0.3037, n = 15-18). However, if we compared % time in open arms in the dim condition to 50% (no preference), we found that female M5 WT showed no difference from 50% (p = 0.2196), compared to M5 HET and M5 KO (ps = 0.0004, 0.0053 respectively) suggesting a subtle deficit in exploration in female M5 KOs under these conditions. Thus, while the deficits in exploration of “riskier” compartments in novel environments tend to be context-specific, the deficit in normal adaptations in behavioral output persist across contexts. Furthermore, this deficit in exploratory behavior appears to be sex dependent.

**Figure 6.**
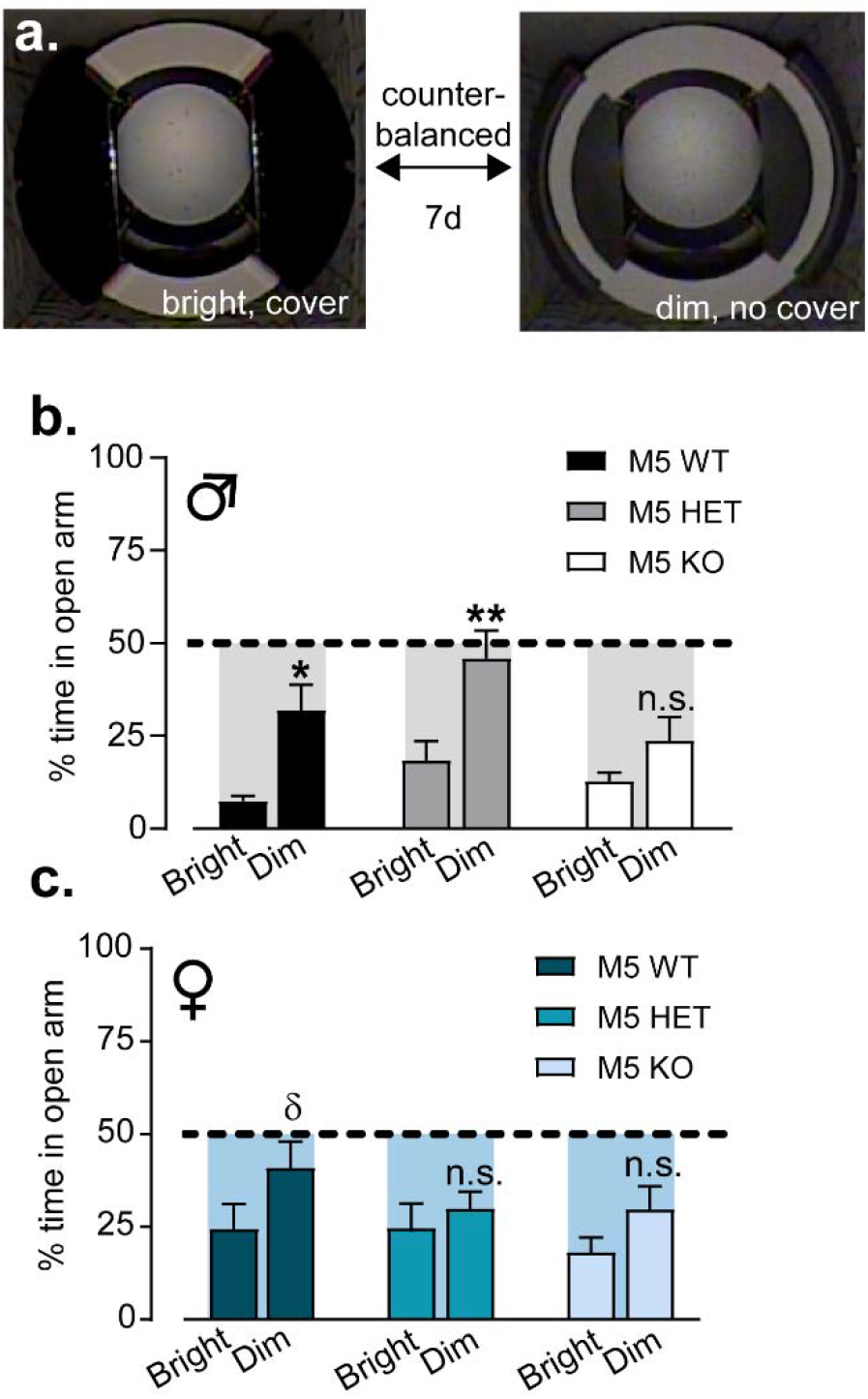
M5 deletion produces deficits in behavioral adaptation in response to changes in environmental conditions in elevated zero maze. a) Representative overhead images of bright, covered (no cover (left) and dim, no cover (right) elevated zero maze conditions. b) % time in open arms for bright and dim conditions for male M5 WT, HET and KO littermates. c) % time in open arms for bright and dim conditions for female M5 WT, HET and KO littermates. Shaded regions highlight the relative difference from 50% (no preference) for each group under each condition.

**Figure 7.**
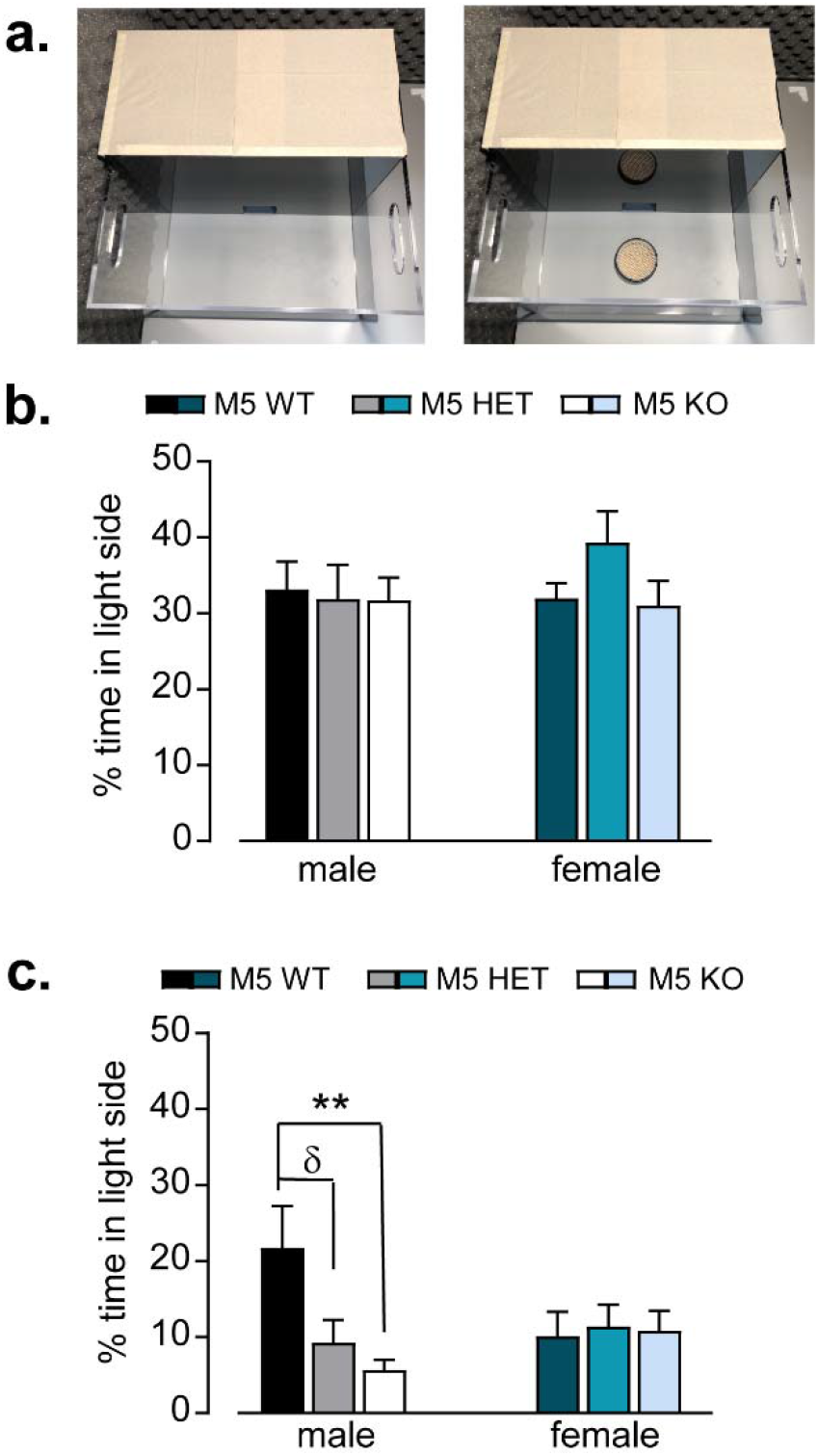
M5 deletion produces sex-dependent selective deficit in novelty exploration. a) a) Representative overhead images of brightly lit light-dark box without (left) and with (right) novel palatable food stimulus. b) % time in light compartment for male and female M5 WT, HET and KO littermates in the absence NF stimulus. b) % time in light compartment for male and female M5 WT, HET and KO littermates in the presence NF stimulus.

### M5 deletion reveals dissociation between generalized anxiety-like behavior and selective deficit in novelty exploration

The discrepancy in phenotype between two classic measures of anxiety-like behavior led us to use yet a third behavioral measure that combines elements of both the OF and EZM – the light-dark box. Following the standard five-minute test, we re-introduced the mice to the chamber with the NF stimuli in the light side. Interestingly, there was no difference in % time in light side between M5 WT, HET and KO for either male or female mice (or between males and females) in the standard five-minute test, similarly to the EZM (2W-ANOVA, F_2,55_ = 0.9393, p = 0.3971, n = 9-13). Novel palatable food placed in the bright light without much habituation was generally aversive across all mice. However, this manipulation revealed a similar aversion to novelty exploration specifically in male mice as shown in OF (2W-ANOVA, sex x genotype interaction, F_2,51_ = 3.325, p = 0.0439; post-hoc Sidak’s t-test, male M5 WT vs. KO, p = 0.007, male M5 WT vs HET, p = 0.0591, n = 8-13). Taken together, this data reveals a behavioral dissociation between general anxiety-like behavior and specific deficits in novelty exploration.

## Discussion

The M5 receptor is an interesting therapeutic target protein because its overall expression is restricted compared to M1-M4, potentially reducing the likelihood of unwanted side effects. In addition, the crystal structure of the M5 receptor has recently been solved, enabling pharmacological targeting of this protein (Vuckovic et al., 2019). Indeed, negative and positive allosteric modulators targeted toward the M5 receptor are actively being developed (Bridges et al., 2010; Bridges et al., 2009; Gentry et al., 2010a; Gentry et al., 2013; Gentry et al., 2010b; Gentry et al., 2010c; Gentry et al., 2014). Manipulations of M5 receptor function have indicated that it may play a role in drug seeking and taking, as well as sensorimotor gating (Basile et al., 2002; Gould et al., 2019; Gunter et al., 2018; Thomsen et al., 2005; Thomsen et al., 2007). However, much remains unknown with respect to more basic characterization of cellular and behavioral functioning for the M5 receptor, with little attention paid to potential sex differences. In this study, we quantify M5 expression in VTA dopamine neurons in males and females and perform a full functional characterization of M5 potentiation of dopamine transmission in both males and females. While there may be some more nuanced differences, overall, we found that both M5 expression and M5 potentiation of dopamine transmission in the NAc were similar in male and female mice. Surprisingly, despite similar cellular function, M5 deletion manifests disparate behavioral deficits in males and females. In females, M5 deletion primarily impairs rearing behavior, a dopamine-dependent motor behavior that is both a form of exploration as well as vigilance. In contrast, M5 deletion in males primarily affected exploratory behavior, particularly of riskier or more exposed areas of our testing chambers or toward novel stimuli. In addition, M5 receptors appear to be important for adapting behaviors in response to new information, a behavioral function that has been attributed to cholinergic interneurons in the striatum. While this study provides a foundation for understanding of M5 function at the cellular, systems and behavioral level, there are many questions that still exist.

### Unique features of M5 modulation of dopamine transmission in the NAc

Our first observation was that there was not an intermediary gene dose effect of OXO-M potentiation of dopamine transmission in the M5 HET mice. This indicates to us that there may be spare receptors on dopamine terminals with the NAc core, which have been shown to exist for several GPCRs including muscarinic receptors (Buchwald, 2019). However, it did seem that both M5 alleles were necessary to trigger the secondary attenuation of the response since a difference in early versus late phase was only detected in the M5 WT and not the M5 HET. The mechanism that mediates this concentration-dependent attenuation is still unclear. It could be that the receptor is desensitized and internalized at higher concentrations. Alternatively, it could be that M2/M4 heteroreceptors, not present on cholinergic interneurons, are recruited at these higher concentrations.

We had expected that increasing concentrations of OXO-M would increase the maximal response as is often the case for other GPCRs. Notably, the majority of research on GPCR regulation of dopamine transmission at the terminals within the NAc has focused on inhibitory modulation of dopamine transmission via Gi/o coupled receptors (i.e. D2, KOR, M2/4) with perhaps the exception of 5-HT2 and mGlu1 receptors (Alex et al., 2005; Britt and McGehee, 2008; Bruton et al., 1999; Shin et al., 2017). In the case of the M5 receptors, increasing concentrations of OXO-M quickened the time to achieve what appears to be a ceiling of maximal potentiation of dopamine release. Increasing concentrations hasten equilibrium within the slice preparation along with receptor binding. We speculate that the coupling effectors that lead to potentiation, likely via Gq-induced release of calcium stores, limit the maximal effect. Furthermore, it is possible that there is a narrow dynamic range of Gq or Gs modulation compared to Gi/o. This an interesting area of future research since the mechanisms of GPCR-mediated modulation of dopamine terminals remain understudied and are still a challenge to probe with our current tools.

Our previous studies demonstrate that repeated stress ablates CRF-mediated potentiation of dopamine transmission at terminals within the NAc core (Lemos et al., 2012). Like CRF, our test using an intermediate concentration of OXO-M showed that repeated stress disrupted M5-dependent potentiation. It is possible that repeated stress elevates ambient levels of acetylcholine to either desensitize and internalize the receptor or recruit inhibitory mechanisms much in the same way high concentrations of OXO-M might evoke the secondary attenuation. It is also possible stress causes reduced M5 receptor expression or uncoupling from effector mechanisms. These will be interesting questions to address in future studies. Our observation that stress disrupts M5 function led us to examine how M5 deletion may lead to anxiety- or depression-related phenotypes.

### The role of M5 receptors in mediating exploratory behaviors

First, we acknowledge the caveat to this work that we are using a constitutive M5 knock-out mouse as opposed to a cell-type specific transgenic and/or viral strategy. Presently, there are no commercially available tools to address this caveat. Our laboratory is developing these tools and we intend to follow-up this study in the future using these novel techniques. This study provides a roadmap for initial validation of subsequent modern techniques.

With this caveat in mind, we explored the consequence of M5 deletion on dopamine-dependent motor, exploratory, and hedonic behaviors. Spontaneous ambulatory behavior was intact. This is consistent with what has been previously reported (Yamada et al., 2001). Yet, we found that in females, M5 deletion disrupted rearing behavior, which is a dopamine-dependent form of vertical inspection and vigilance displayed in novel environments. In contrast, M5 deletion manifested differently in males. Males displayed disruption in exploration of the center of a novel open field as well as disruption in the exploration of novel appetitive stimuli. For both males and females, these disruptions in exploratory behaviors were not accompanied by anhedonia assayed using the sucrose preference test, despite evidence that both scopolamine and reduction in cholinergic interneuron firing lead to reduction in sucrose seeking and preference (Addy et al., 2015; Cheng et al., 2019; Navarria et al., 2015; Warner-Schmidt et al., 2012). These data indicate that cholinergic modulation of hedonia is dependent on activation of M1,2,3 or 4. Both novelty and risk preference have been linked with increased drug taking and seeking (Belin et al., 2011; Wingo et al., 2016). Therefore, it is consistent with the literature that a disruption in novelty exploration and risk aversion would occur in the same genotype in which decreases in drug seeking and taking are frequently observed.

Our findings are not consistent with a recent study done by the Addy laboratory in rats in which rats received intra-VTA infusion of physostigmine with or without co-administration of the M5 negative allosteric modulator (NAM), ML375 (VU6000181) (Nunes et al., 2019). The investigators found that intra-VTA physostigmine, a treatment that would elevate ACh levels in the VTA, produced anhedonia and anxiety-like behavior in both males and females assayed by the sucrose preference test, EPM and forced swim test, respectively. Interestingly, co-infusion of the M5 NAM prevented these effects only in males. This sex-dependent component is consistent with the sexual dimorphism we have observed. However, this study suggests that M5 function in the VTA produces aversion and is anxiogenic. There may be several reasons for the inconsistencies between our findings and those of Nunes et al., including species differences, developmental compensation due to constitutive M5 deletion, regional specificity of the Nunes et al. manipulations, potential non-selectivity of the ML375 compound and nuances in the execution of our behavioral paradigms. For example, we used 4% sucrose, while they used 1% sucrose. We believe that some of these inconsistencies may be reconciled with the development of cell-specific transgenic and viral techniques.

### The role of M5 receptors in behavioral adaptation

In the open field test, control mice show a reduction in center exploration of the novel open field over time. However, this habituation is not present in mice with constitutive M5 deletion. Likewise, in the EZM, mice adjust their time spent in the open compartments based on the lighting conditions and the level of enclosure of the closed compartments. This is disrupted in both male and female M5 KO mice. Work from the Cain lab has demonstrated that M5KO mice have a deficit in prepulse inhibition (Thomsen et al., 2007). Though prepulse inhibition is considered a sensorimotor gating assay and animal model for features of schizophrenia, at a broader level it relates to the ability to integrate information and adjust the behavioral response. Taken together, it is possible that M5 receptors play a broader role in behavioral updating in response to changes in environmental contingency. Indeed, it has been suggested that striatal cholinergic interneurons are important for behavioral updating and flexibility (Apicella, 2017; Bradfield et al., 2013). It is plausible that M5 is a downstream effector of cholinergic interneuron responses to environmental changes.

## Conclusion

There are several open questions remaining that are beyond the scope of this initial study that we hope are actively pursued by other laboratories in addition to our own. Considering both the therapeutic potential and interesting biological function of the M5 receptor, we hope this study brings M5 to the forefront of GPCR and neuroscience research.

## Author Contributions

Conceptualization: J.C.L. and V.A.A., Methodology: J.C.L, J.A.R., S.M.L.F., V.A.A. Investigation, validation, and analysis: J.C.L., J.A.R, S.M.L.F., L.C.W., S.A.M. Writing manuscript: J.C.L., Manuscript editing: J.A.R., S.M.F., L.W., V.A.A., J.C.L Funding: J.C.L., Resources: J.C.L. Supervision: J.C.L.

## Acknowledgements

This work was supported by MH109627 (JCL), MH122749 (JCL), Department of Neuroscience and Medical Discovery Team on Addiction start-up funds (JCL) and AA000421 (VAA). We would like to thank Dr. Jürgen Wess at NIDDK/NIH for providing the M5 KO mouse line. We would like to thank Ms. Rachel Dick for help with cryosectioning used for RNAscope techniques. We would like to thank Dr. Sade Spencer and the Spencer lab for performing pilot western blot experiments. We would also like to thank Drs. Erin Calipari, Mark Thomas, Michael Bruchas, Larry Zweifel, Anna Ingebretson, Chris Petersen and Ms. Elizabeth Souter for their helpful input and valuable comments in the synthesis of this work.

## Extended Figures and Figure Legends

**Extended Figure 1.**
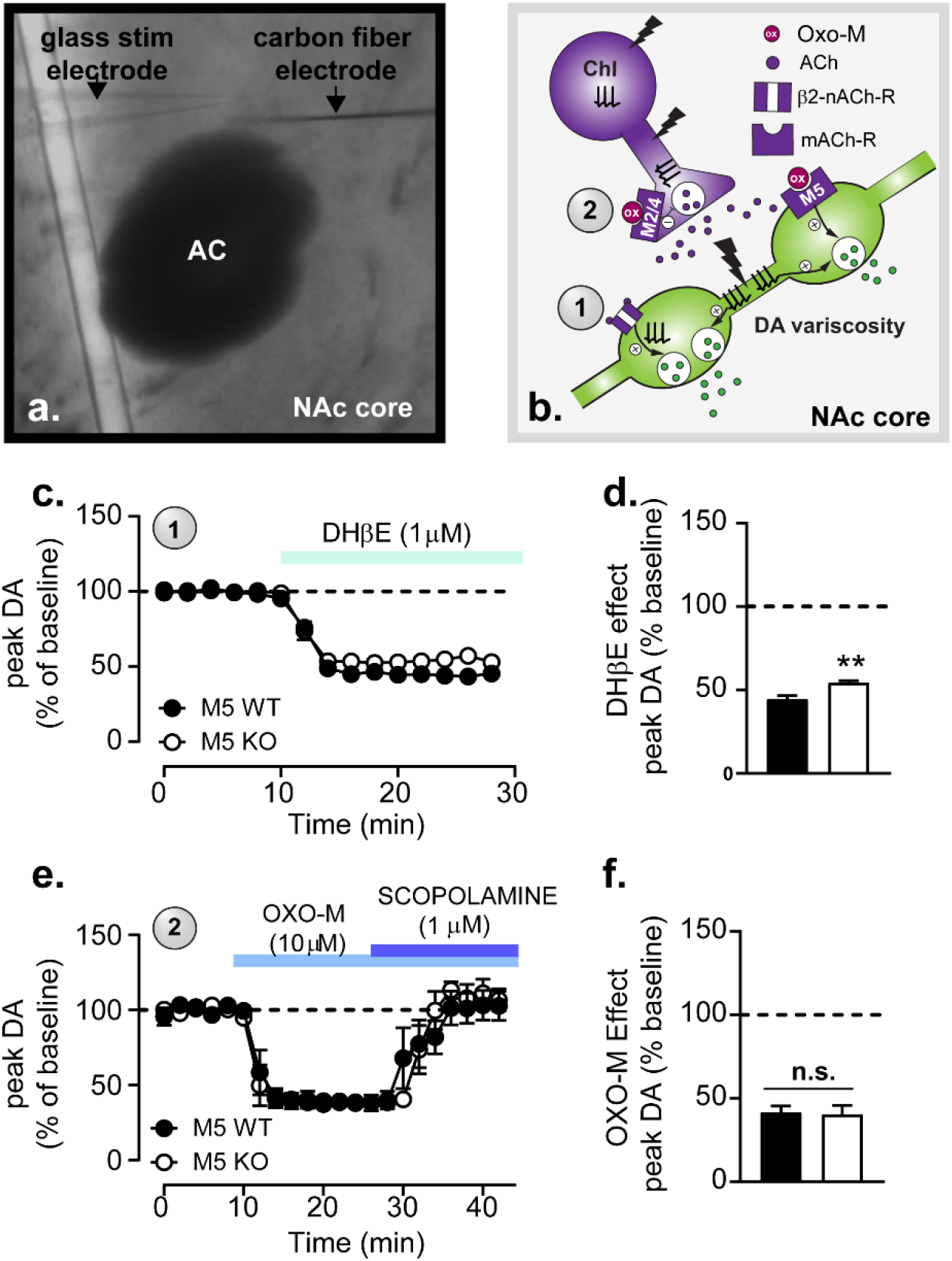
related to Figure 1. β2-nACh-R and M2/M4 autoreceptor function are intact following M5 deletion. a) Low power DIC-IR image demonstrating stimulating electrode (left) and carbon fiber electrode (right) placement within the NAc core. b) Schematic diagram depicting cholinergic and dopaminergic interactions that are engaged during the voltammetry paradigm used to execute the experiments in c-f. c) Time course of the normalized peak amplitude of electrically evoked dopamine transients prior to and following bath application of DhβE (1μM) in NAc slices taken from M5 WT and KO mice. d) Mean steady state effect of DhβE application on dopamine transients recorded from M5 WT and M5 KO mice e) Time course of the normalized peak amplitude of electrically evoked dopamine transients prior to and following bath application of OXO-M (10μM) in NAc slices taken from M5 WT and KO mice with no DhβE onboard. f) Mean steady state effect of OXO-M application on dopamine transients recorded from M5WT and M5KO mice in slices maintained in ACSF with no DhβE onboard.

**Extended Figure 2.**
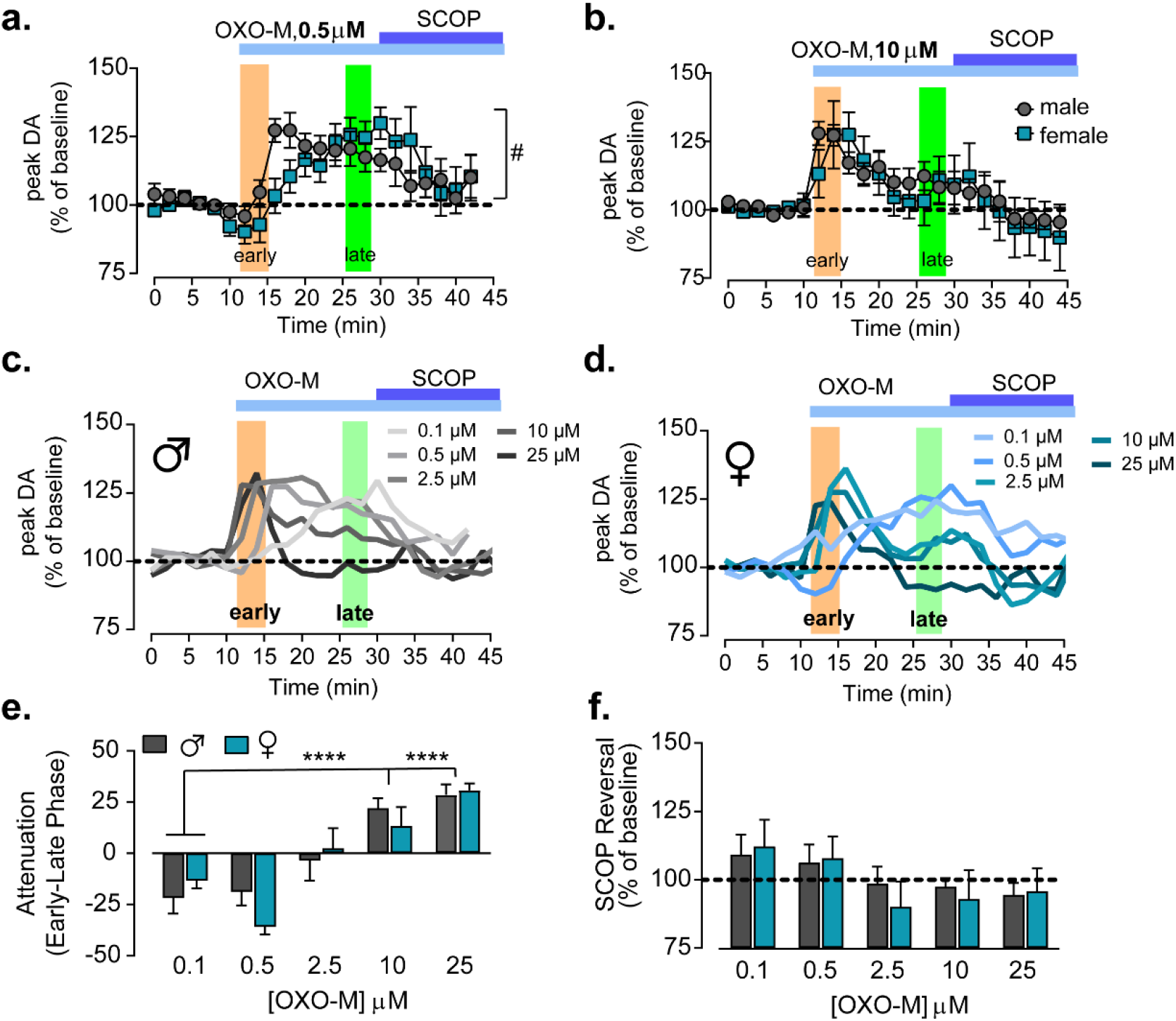
related to Figure 3. The effect of 0.5 μM and 10 μM OXO-M on dopamine transmission in male and female mice. a) Time course of normalized peak amplitude of electrical evoked dopamine transients recorded from *ex vivo* NAc slices taken from male (grey) and female (teal) M5 WT mice in which OXO-M (0.5 μM) followed by scopolamine (1μM) is bath applied to the slice. There was a significant sex x time interaction. (2W ANOVA, sex x time interaction, F_21,229_ = 1.879, p = 0.0133, n = 6-7) b) Time course of normalized peak amplitude of electrical evoked dopamine transients recorded from *ex vivo* NAc slices taken from male (grey) and female (teal) M5 WT mice in which OXO-M (10 μM) followed by scopolamine (1μM) is bath applied to the slice (n = 7-22). c,d) Overlaid time courses for OXO-M (0.1, 0.5, 2.5,10, 25 μM) for male (c) and female (d) M5 WT mice. e) Attenuation (% baseline, late phase - % baseline, early phase) observed during OXO-M bath application (0.1, 0.5, 2.5,10, 25 μM) for male and female mice. (2W-ANOVA, main effect of concentration, F_4,68_ = 21.15, p < 0.0001, post-hoc t-test, 0.1 vs. 10 μM, p < 0.0001, 0.01 vs. 25 μM, p < 0.0001). f) Scopolamine reversal (%baseline) observed during OXO-M + Scopolamine bath application. There were no significant main effects of sex or concentration nor was there was there a significant sex x concentration interaction.

